# Community, Belonging, and Peer-Engagement: Participant Experiences Reflect Consistency of Facilitation Practices in Inclusive STEM Teaching Learning Communities

**DOI:** 10.1101/2025.01.13.632844

**Authors:** Diane Codding, Alexandria H. Yen, Haley Lewis, Vanessa Johnson-Ojeda, Sarah Chobot Hokanson, Bennett B. Goldberg

**Affiliations:** Northwestern University; Boston University; University of Utah

**Keywords:** teaching, learning community, STEM education, faculty development, positive peer engagement

## Abstract

A primary reason for the persistent underrepresentation of minoritized people in STEM careers are their marginalizing experiences in STEM courses, which has spurred efforts to prepare STEM faculty to implement inclusive teaching practices. The Inclusive STEM Teaching Project (ISTP) addresses this nationwide challenge by providing STEM faculty a free, online course and associated learning communities (LCs) led by project-trained facilitators. ISTP centers identity, power, privilege and positionality and has successfully shifted educators’ mindsets and abilities. A central question is whether local LCs across contexts maintain equity-minded core practices and achieve inclusive, reflective practitioner outcomes. Our mixed methods analysis of LC participant (*n* = 165) and facilitator (*n* = 83) matching survey data spanning five course offerings over two years directly explores the relationship between the learning environments created by facilitators and their participant experiences. Findings demonstrate high fidelity of implementation to training by facilitators that resulted in consistent LC experiences of participants, especially around building a community, sense of belonging and positive peer engagement. Participants overall speak to the importance of facilitated LCs as an inclusive space where they can critically engage with one another as they reflect on, discuss, and ultimately improve their inclusive teaching practices.

---

Each year, historically marginalized students leave science, technology, engineering, and math (STEM) majors at a disproportionately higher rate than their majority peers (Riegle-Crumb, 2019; Thiry, 2019). Research has shown female and racially minoritized students often leave due to an alienating climate and exclusionary practices that contribute to students experiencing a lack of belonging in the field (Handelsman *et al*., 2022). In acknowledgement of this persistent inequity, there has been a national drive to prepare STEM faculty to implement inclusive teaching practices to address the underrepresentation of historically marginalized students (see *Broadening Participation in STEM*, n.d.). Inclusive teaching has been shown to reduce STEM performance gaps (Handelsman *et al*., 2022), encourage student persistence (Riegle-Crumb, 2019; Thiry, 2019), and increase the participation of historically excluded groups in STEM economy and innovation (NCSES, 2023). Yet, STEM faculty receive limited professional development in pedagogy and instruction, and even less still in inclusive teaching (Addy *et al*., 2021). Positive change has been achieved in local, institutional efforts in inclusive teaching (Elliot *et al*., 2016; Macaluso *et al*., 2020; O’Leary *et al*., 2020) and a few cross-institutional projects have been successful (Esquibel *et al*., 2023; Gehrke and Kezar, 2015; Pelletreau *et al*., 2018). These initiatives largely address inclusivity indirectly through active learning and student-centered pedagogy; our professional development engages faculty in critical self-reflective work to advance diversity, equity, and inclusion (DEI) in STEM education on a national scale.

The Inclusive STEM Teaching Project (ISTP), the focus of this paper, addresses this challenge by providing targeted professional learning for STEM faculty that centers identity, power, privilege, and positionality. Building on an introspective foundation, the ISTP has successfully shifted educators’ mindsets and abilities through an online course paired with optional learning communities (LCs) and affinity groups led by project-trained facilitators (Calkins *et al*., 2024; Codding *et al*., 2024). This paper examines these course-aligned LCs as a key professional development methodology, addressing the question as to whether local LCs across contexts maintain equity-minded core practices and achieve inclusive, reflective practitioner outcomes. Specifically, this paper examines the impact of ISTP LCs on participant experiences, learning outcomes, and intentions to implement inclusive teaching practices. In addition, by analyzing LC participant (*n* = 165) and facilitator (*n* = 83) survey data spanning five course offerings, 55 LCs, institutional context, and diverse demographics, we were able to directly explore the relationship between participant experiences and the learning environments created by facilitators. This mixed methods inquiry was guided by the following research questions:

1. What learning environment are participants experiencing within LCs?
2. What are the relationships between LC facilitators’ inclusive learning facilitation and participant experiences?
3. How are participants planning to implement inclusive teaching practices following LC participation?

Findings demonstrate high fidelity of implementation to training by facilitators that resulted in consistent LC experiences among participants, especially around building community, sense of belonging, and positive peer engagement. Strong matching of facilitator actions and participant experiences confirm inclusive and effective LCs, while areas of consistent difference of delivery and experience are likely due to reasonable differences in expectations. We also note, surprisingly, that our results are agnostic to individuals’ identities or institutional context.

Participants overall speak to the importance of facilitated LCs as an inclusive space where they can critically engage with one another as they reflect on, discuss, and ultimately improve their inclusive teaching practices. Grounded in the online course and framed by effective, inclusive facilitation, participants turned toward each other to share experiences, collaboratively problem solve, and develop concrete strategies for implementing inclusive teaching practices in their classrooms.

## Inclusive STEM Teaching Project

The ISTP is a national professional development initiative designed to advance equitable STEM education by engaging faculty, staff, postdoctoral scholars, and doctoral students in developing the knowledge, skills, and beliefs necessary for inclusive teaching. The project offers an asynchronous online course that engages participants in critical self-reflection that centers identity, power, and privilege while upholding the principle of “do no harm” (Calkins *et al*., 2024). The ISTP project explicitly sought to ‘do no harm’ through carefully curating curriculum, learning environments, and facilitated discussions to avoid putting minoritized individuals in marginalizing or re-traumatizing situations (Rhodes *et al*., 2019; Tajima, 2021). Course participants progress through six linked modules: introduction and framing; diversity, equity and inclusion in learning and teaching; instructor identity; student identity; creating an inclusive STEM course; and climate in the STEM classroom. At the time of publication, ISTP has run seven free open online courses with 12,719 participants, 2,872 (23%) of whom completed the full online course. Completion rates for those who answered one question (68%) or visited one page (56%) are five times higher than the average for free online courses (Jordan, 2015; Reich, *et al*., 2019).

Course participants also have the option to participate in locally run, synchronous LCs led by project-trained facilitators. To date, ISTP has trained 513 facilitators in 164 institutions, including 2-year technical and community colleges, 4-year predominantly undergraduate and liberal arts schools, comprehensive universities, and research intensive institutions. LCs typically meet weekly for 60-90 minutes over the 6-week course run, during which they connect with others in their local community, engage with course content, and contextualize applications of their learning. To ensure effective facilitation, ISTP uses a model for selecting, training, and supporting facilitators that utilizes a co-facilitation model and prioritizes prior DEI experience (Codding *et al*., 2024). Specifically, ISTP (1) selects teams of motivated facilitators with an average of seven years of prior DEI experience, (2) trains facilitator teams in a succinct training model that focuses on normalizing facilitator practices, and (3) supports facilitators with a carefully curated workbook and synchronous drop-in community discussions. This training model leverages LC facilitators’ existing skillsets while also providing training that research has shown to increase facilitator confidence significantly in four key areas: facilitating DEI conversations, creating open dialogue, leading conversations centered on identity, and managing difficult moments in DEI conversations (Codding *et al*., 2024). This paper builds on our previous work by examining participants’ learning and experiences in comparison to how facilitators’ self-report cultivating inclusive LCs.

STEM mentoring practice has been greatly advanced in higher education nationwide through similar large-scale, train-the-trainer (facilitator) models that implement local LCs with a common curriculum (Pfund *et al*., 2017; Rogers *et al*., 2018). Studies of these efforts have demonstrated that their multi-day, in-person training is effective in educating facilitators, and distal research has shown the high confidence, effective delivery and program growth as reported by trained facilitators across multiple institutions. Our study adds to this literature by going further and directly exploring the relationship between participant experiences and the learning environments created by facilitators.

## Faculty Learning Communities

Faculty LCs have been shown to help instructors develop confidence for implementing and sustaining new pedagogical practices (Cox, 2004; Furco and Moley, 2016; Gehrke and Kezar, 2016; Nadelson *et al*., 2013; Price *et al*., 2021). Additionally, faculty who participate in inclusive STEM teaching LCs have been shown to develop greater long-term interest in using new pedagogical strategies (Esquibel, 2023; Nadelson *et al*., 2013; Tinnell *et al*., 2019) and their students report feeling a greater sense of engagement, inclusion, and encouragement (Elliot *et al*., 2016; Jaimes *et al*., 2024). LCs can foster long-term commitment to inclusive teaching by promoting community and collaboration among participating faculty (Cherrington et al., 2018; Pelletreau *et al*., 2018), increasing commitment to student-centered teaching (Anderson and Finelli, 2014; Nadelson *et al*., 2013), and generating interest in learning about student identities (Macaluso, 2020). LCs provide a supportive space for critical reflection and lead instructors with marginalized identities to report feeling a greater sense of belonging within their institution (Drane *et al*., 2019; O’Meara *et al*., 2019). Faculty LCs that focus on community cultural wealth help faculty identify teaching strategies and curriculum that promote a strong sense of belonging in STEM classrooms by emphasizing students’ cultural identities and histories (Cavazos *et al*., 2024). Recent studies have also shown that institution-specific LCs are an important and effective space for reflecting on instructor and student identities (Cavazos *et al*., 2024; Drane *et al*., 2019; O’Meara *et al*., 2019). ISTP furthers institution-specific efforts to advance inclusive teaching and enact equitable change by adopting a structure that pairs our online course with LCs designed to be implemented locally while also targeting a diverse range of institutions and teaching environments at a nationwide scale.

## Online Course Associated Learning Communities

Pairing an online course with facilitated LCs has been shown to be an effective approach for professional development in pedagogy (Goldberg *et al*., 2023; Sun *et al*., 2023). Research has shown that opportunities for participant interactions directly influence the successful retention of learners (Alemayehu and Chen, 2021), and LCs provide a facilitated space for critical peer engagement (Codding *et al*., 2024). When implemented alongside online courses, LCs create a space where participants can share learning strategies and engage with differing opinions (Chen and Chen, 2015). In the case of ISTP, the online course builds a common foundation of inclusive teaching approaches specific to STEM, which participants augment by engaging in critical and often difficult discussions during facilitated LCs. ISTP specifically focuses on helping instructors recognize and describe how identity, positionality, and power (of both students and instructors) influences STEM retention and belonging in their own classrooms (Sue *et al*., 2009; Yosso, 2005). The online course offers moderated discussion boards and optional affinity groups that meet synchronously three times each course run, but the majority of the course is self-paced (Calkins *et al*., 2024). For example, the “My Inclusive Framework” includes a series of questions that allow participants to individually reflect on course content and application. In comparison, the LCs utilize a group learning modality—a community-focused discussion format that supports participants as they reflect on and share some of the challenges of implementing culturally relevant and sustaining pedagogy, such as competing responsibilities, little training, and lack of reward (Anderson *et al*., 2011; Brownell and Tanner, 2012; Shadle *et al*., 2017).

## Method

### Data Collection

Data for this mixed-methods study were collected over five course runs (summer 2021, fall 2021, spring 2022, fall 2022, and spring 2023) under approval from the Northwestern University Institutional Review Board (STU00207792) with reciprocal agreements at partnering institutions. Surveys were distributed via Qualtrics to all LC participants and active LC facilitators following each course run. The Participant Survey (Supplemental Material) included 30 questions that address participant experiences during LC meetings, peer interactions, and inclusive teaching outcomes. The Facilitator Survey (Supplemental Material) included 48 questions that address facilitation methods and pedagogy, perceived participant experiences, similarity and difference in general DEI facilitation, and utilization of various facilitation resources. Both surveys included a mix of 6-point Likert scale, open response, sliding scale, and multiple choice questions. Survey questions for both LC facilitators and participants included a series of similar questions, some identical and some not, asking facilitators to reflect on the LC environment they created and asking participants to reflect on how they experienced the LC environment. Questions from the two surveys were aligned closely enough to allow for a visual comparison of Likert-scale (see Supplemental Material for a table of aligned survey questions).

### Quantitative Data Analysis

The quantitative data analysis was run using tidyverse (Version 2.0.0; Wickham *et al*., 2019), ggpubr (Version 0.6.0; Kassambara, 2023a), and rstatix (Version 0.7.2; Kassambara, 2023b) packages in R (Version 4.2.2; R Core Team, 2023). All analyses were performed on de-identified data. Years of prior DEI experience and prior experience as DEI participants and/or facilitators in DEI and faculty LCs were compared between participants and facilitators. Paired *t tests* were used to determine if there were significant differences between the faculty/non-faculty, white/non-white, and women/men demographic groups as it relates to (a) LC participation, (b) LC facilitator evaluation, (c) post LC course impressions, and (d) LC social identity. These demographic pairings were chosen because they each provided enough statistical power (*n* > 50) for response comparisons.

Normalized histograms were created to compare the distribution of LC participant and facilitator responses to aligned questions (see Supplemental Material). As regression is possible only for the few exact matching questions, we have chosen to instead plot the overall distributions of facilitator and participant responses. The exact matching questions allow us to benchmark the level of correspondence and thus we argue that it is reasonable to compare the distribution of responses for not only the exact matching questions but closely aligned questions as well. For exact matching questions, we calculated the distribution of the Likert-scale distance between participant and facilitator responses for overlapping LCs, which measures the percentage of exact, one scale point difference and two scale point difference in cohort responses. We chose to down-select when multiple participant responses for a given LC were present using the median among participants, which is a good measure based on the narrow response distribution. We can apply this procedure only for exactly matching questions (e.g., establishing community norms) and, as demonstrated in the results section below, find a high level of congruence between the distributions of all responses that we plot and the distributions when the subset is limited to overlapping LCs and exact question correspondence.

### Qualitative Data Analysis

Qualitative data were organized into course runs and analyzed using Dedoose (v.9.0.17). Our qualitative codebook (Supplemental Material) was collaboratively developed using a grounded theory approach (laser and Strauss, 1967) by two researchers who completed two rounds of independent coding and collaboratively reached a consensus. The final thematic codebook included five categories: identity and awareness, inclusive community, LC group dynamics, discussion approaches, and teaching and pedagogy. Open-ended responses were coded holistically within the context of each survey question, which resulted in the application of a single code unless multiple examples were specified. Responses related to planned changes in teaching were also coded as either *tangible* (i.e., specific, concrete strategies) or *aspirational* (i.e., descriptions of outcomes that respondents hope to achieve). Qualitative data were analyzed by the co-first authors, both of whom identify as women scholars from majority identities in STEM (white and East Asian respectively). We calculated inter-rater reliability (IRR) in Dedoose using a code application test on 20% of the data, which allowed us to generate a pooled kappa to summarize interrater agreement across multiple codes (Cohen, 1960; de Vries *et al*., 2008). The results of the IRR indicated a pooled kappa (κ) of 0.62, meeting the standard for *good agreement* (Cicchetti, 1994; Fleiss, 1971; Landis and Koch, 1977). After reviewing the questions, with κ ≥ 0.60, the two researchers adjusted the codebook and coded the remaining data.

### Participants

We invited all LC participants (*n* = 775) and active facilitators (*n* = 231) following the completion of each of the five course runs, of whom 280 participants and 117 facilitators completed the survey (response rates of 36.1% and 50.6% respectively). We excluded 115 participant surveys and 34 facilitator surveys for either not providing research consent or completing less than 50% of the survey. The cleaned data set included responses from repeat facilitators (*n* = 10) who indicated that their most recent facilitation experience was sufficiently different from prior experiences, so each represented a unique data point. Distinct IDs were assigned to each of the remaining participants (*n* = 165) and facilitators^1^ (*n* = 83), who represent a total of 56 institutions.

Table 1 presents demographic data for LC participants and facilitators, including gender, race and ethnicity, institution type, academic position, academic discipline, and prior DEI experience. Facilitators applied and were accepted into the program based partially on prior DEI experience (M = 7.69, SD = 5.82), so it is interesting to note that participants, who self-selected into LCs, reported similarly high levels of DEI experience (M = 5.94, SD = 5.39).

**Table 1.** Participant Demographics.

| Factor | LC Participants |  | LC Facilitators |  |
| --- | --- | --- | --- | --- |
|  | *n* | % | *n* | % |
| **Gender** |  |  |  |  |
| Woman | 99 | 60.0% | 60 | 72.3% |
| Man | 52 | 31.5% | 17 | 20.5% |
| Gender-queer/Nonbinary | 6 | 3.6% | 1 | 1.2% |
| Trans Woman | 2 | 1.2% | 0 | 0.0% |
| Trans Man | 0 | 0.0% | 0 | 0.0% |
| Prefer not to respond/No response | 6 | 3.6% | 5 | 6.0% |
| **Race/Ethnicity** |  |  |  |  |
| White | 115 | 69.7% | 59 | 71.1% |
| Asian/Asian American & Pacific Islander | 27 | 16.4% | 9 | 10.8% |
| Latino/a/e/x/Hispanic | 16 | 9.7% | 5 | 6.0% |
| Black/African American | 8 | 4.8% | 8 | 9.6% |
| Middle Eastern/North African | 5 | 3.0% | 2 | 2.4% |
| Alaskan Native/Indigenous | 1 | 0.6% | 1 | 1.2% |
| Multiracial | 12 | 7.3% | 10 | 12.0% |
| Prefer not to respond | 3 | 1.8% | 3 | 3.6% |
| **Institution Type** |  |  |  |  |
| Research University | 102 | 61.8% | 57 | 68.7% |
| Comprehensive/Regional University | 23 | 13.9% | 12 | 14.5% |
| Liberal Arts college | 19 | 11.5% | 4 | 4.8% |
| Community College/2-year Institution | 13 | 7.9% | 8 | 9.6% |
| Technical University | 2 | 1.2% | 2 | 2.4% |
| Other | 3 | 1.8% | 2 | 2.4% |

<sup>1</sup> Facilitator data from the first four course runs ( $n = 71$ ) were analyzed and reported on in a previous publication (Coddling *et al.*, 2024).

Table 1 continued.
| Factor | LC Participants |  | LC Facilitators |  |
| --- | --- | --- | --- | --- |
|  | *n* | % | *n* | % |
| **Academic Position** |  |  |  |  |
| Faculty | 100 | 60.6% | 46 | 55.4% |
| Postdoctoral Scholar | 20 | 12.1% | 2 | 2.4% |
| Graduate Student | 18 | 10.9% | 3 | 3.6% |
| Staff | 17 | 10.3% | 28 | 33.7% |
| Other | 10 | 6.1% | 4 | 4.8% |
| **Academic Discipline** |  |  |  |  |
| STEM | 137 | 83.0% | 59 | 71.1% |
| Non-STEM | 28 | 17.0% | 24 | 28.9% |
| **Prior DEI Experience** |  |  |  |  |
| Attended DEI Events | 145 | 87.9% | 80 | 96.4% |
| Facilitated DEI Events | 37 | 22.4% | 68 | 81.9% |
| Organized DEI Events | 43 | 26.1% | 60 | 72.3% |

Table 2 presents details of the institutional overlap present in our data set. Of the 56 institutions represented in the study, we have overlapping facilitator and participant responses from the 34 institutions. Additionally, we were able to break down the data further to reveal the overlapping responses from participants and facilitators in the same LC (*n* = 43) based on their corresponding institution and course run. Thus, we can state with a high level of confidence that our participant and facilitator responses represent self-reflections from a representative cross section of individuals who shared experiences in the same LCs.

**Table 2.** Institutional and Learning Community Alignment.

|  | LC Participants |  | LC Facilitators |  |
| --- | --- | --- | --- | --- |
|  | *n* | % | *n* | % |
| Overlapping Institutions ($n = 34$) | 138 | 83.64% | 71 | 85.54% |
| Overlapping Learning Communities ($n = 43$) | 105 | 63.64% | 60 | 72.29% |

*Note.* Overlapping institutions indicate institutions from which we have both participant and facilitator responses, while overlapping LCs indicate responses from corresponding institutions and course runs.

## Results

### What learning environment are participants experiencing within LCs?

#### Participants Report Positive and Productive Spaces for Peer Engagement

Participants self-reported generally high agreement regarding a range of experiences in ISTP LCs (see Figure 1; Supplemental Material). Participants reported that community norms were established at the beginning of LCs (M=5.49, SD=0.93), and that they actively upheld these norms (M=5.58, SD=0.74). Regarding LC facilitators, participants strongly agreed that facilitators were responsive to making in-the-moment adjustments to the LC based on feedback (M1=5.44, SD=0.89) and actively modeled inclusive practices in STEM teaching (M=5.49, SD=0.85). Reflecting on their own engagement, participants strongly agreed that they had regularly attended their LC sessions (M1=5.55, SD=0.84).

**Figure 1.**
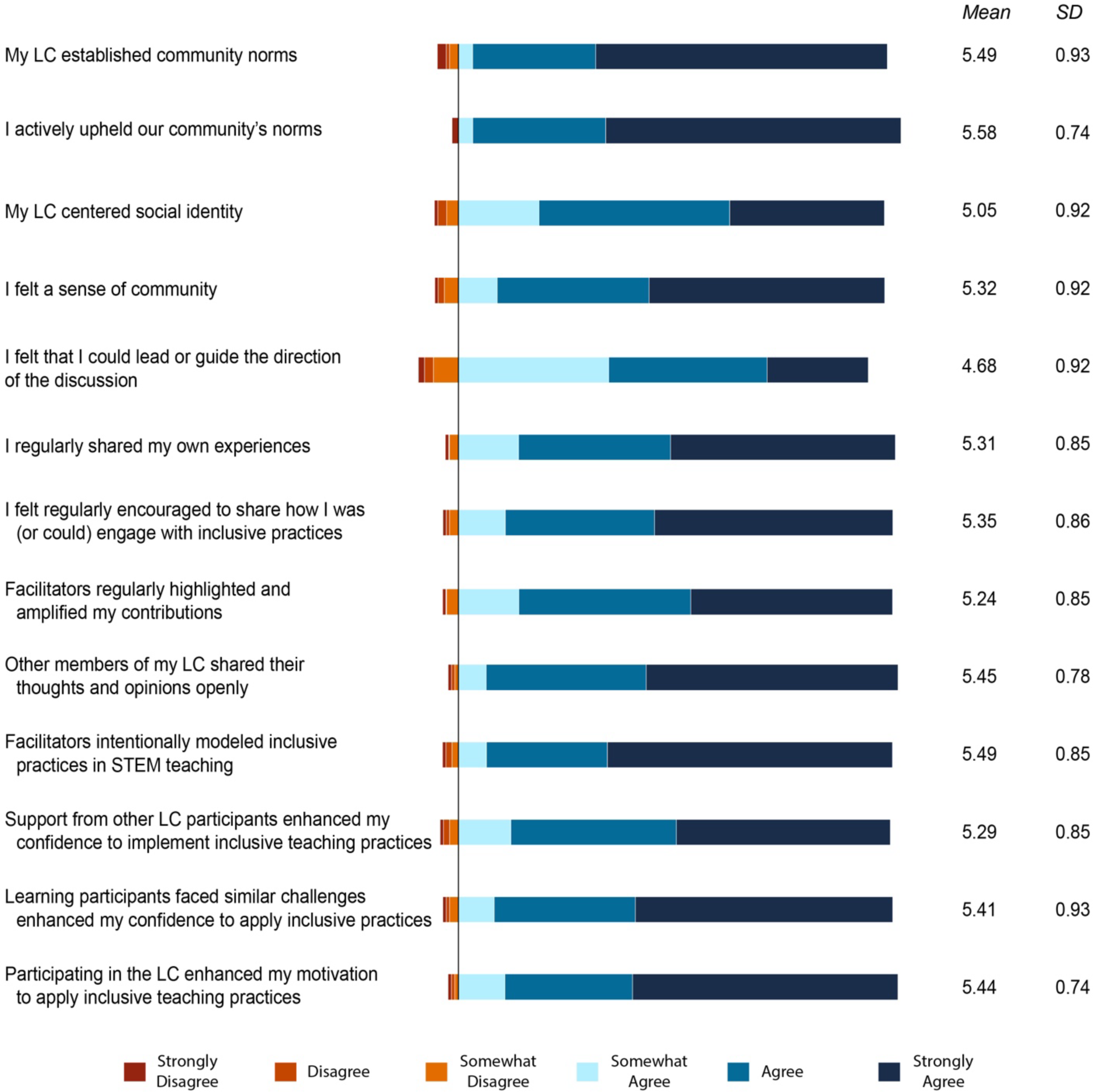
Participants Report Positive Experiences in ISTP Facilitated Learning Communities *Note.* Table of corresponding data can be found in the Supplemental Material

Finally, participants strongly agreed that “the opportunity to hear that other participants faced the same challenges I do enhanced my confidence to apply inclusive teaching practices” (M=5.41, SD=0.93). Participants also agreed that “participating in the LC has enhanced my motivation to apply inclusive teaching practices in my own instruction” (M=5.44, SD=0.74). Participants reported feeling a strong sense of community within their LCs (M=5.32, SD=0.92). However, they were slightly less confident in their ability to implement inclusive teaching following LC participation (M=4.98, SD=1.15).

Results show that, of the 66 paired *t*-tests run, there were only five instances of significant differences among the three demographic pairings (i.e., white/non-white, faculty/non-faculty, women/men) for any of the four measures: (a) LC participation, (b) LC facilitator evaluation, (c) post LC course impressions, and (d) LC social identity (Table 3).

**Table 3.**
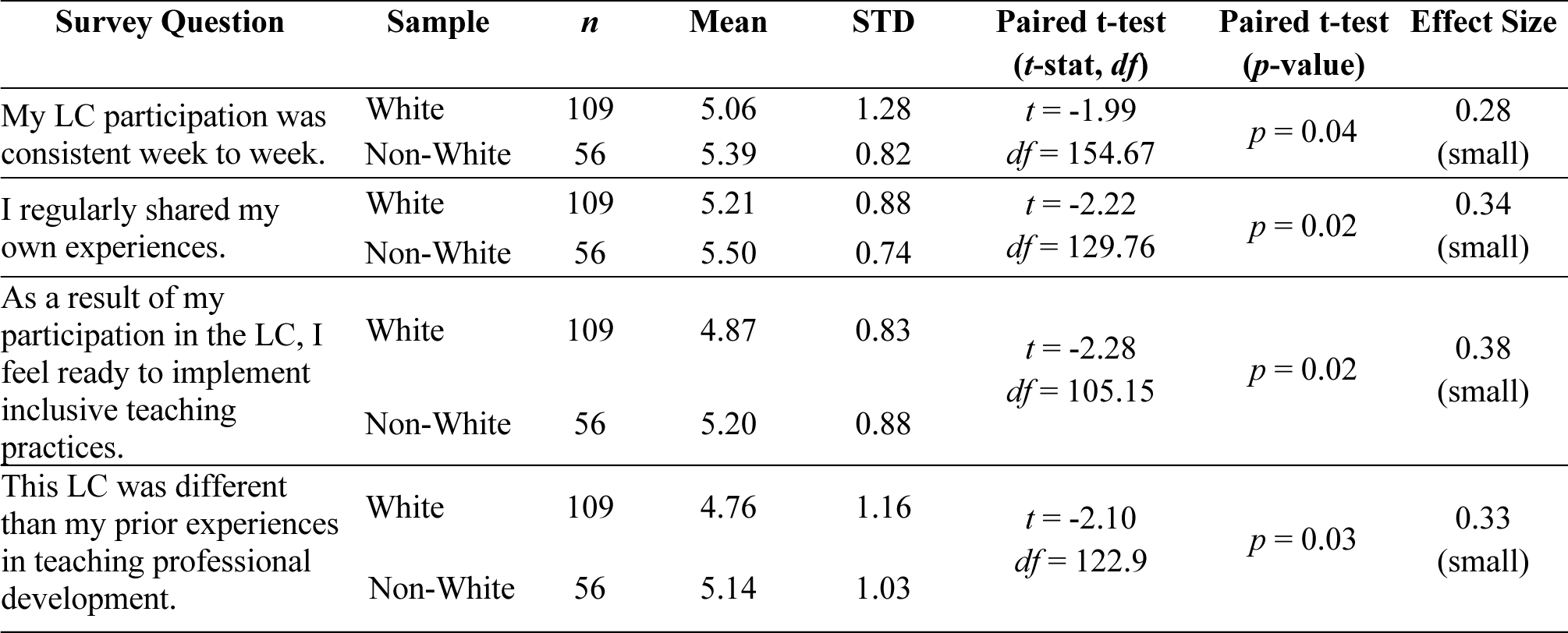
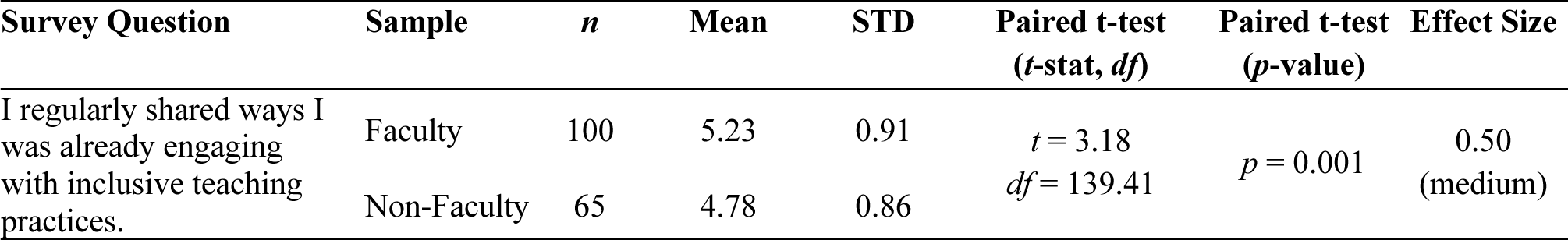
Statistically significant differences among demographic pairings (5 of 66 *t*-tests run)

There were significant differences in the agreement between white and non-white participants for the questions regarding LC participation and post LC course impressions. For LC participation, non-white participants tended to have a higher average agreement that their LC participation was consistent from week to week (M1: 5.06, M2: 5.39). Similarly, non-white participants tended to share more of their personal experiences when compared to white participants (M1: 5.21, M2: 5.50). For the post LC course impressions measure, non-white participants felt more ready to implement inclusive teaching practices after participating in a LC than white participants (M1: 4.87, M2: 5.20) and they also tended to have higher agreement that the LC was different than prior experiences in teaching professional development (M1: 4.77, M2: 5.14). There were also significant differences between faculty and non-faculty agreement for the LC facilitator evaluation measure. Participants with faculty positions tended to agree that they regularly shared ways that they were already engaging with inclusive teaching practices more than non-faculty participants (M1: 5.23, M2: 4.78). Nonetheless, in all of these instances, the significant differences were either small or medium in effect size (Cohen’s d |0.8|) which suggests that there were no substantial differences between participants’ responses overall.

#### Participants Report a Strong Sense of Community

Qualitative results reveal a strong sense of community among LC participants. When asked to share examples of why they experienced a sense of community, participants emphasized having the opportunity to share experiences, finding shared commonalities with their peers, and experiencing productive interactions. While participants mentioned facilitators more frequently while discussing shared experiences, their responses primarily focused on their interactions with LC peers.

##### A Facilitated Space for Sharing

Participants broadly felt that the LC was an ideal space to share their own experiences. Participants often credited facilitators with constructing learning activities and creating an LC structure that allowed for opportunities to share. As one participant explained, “The facilitator was just a fantastic person and great at providing structure for conversations while also encouraging others to speak.” Participants cited “small group work” and working in pairs as a way in which they felt comfortable sharing and learning more about their peers. Some participants attributed these spaces to facilitators, while others did not. As one participant shared, “I usually get overwhelmed with learning names, but with the small groups and Zoom it was much easier to learn names and feel a sense of community.” Participants also provided concrete descriptions of how facilitators applied (i.e., modeled) inclusive teaching strategies to encourage engagement. “The facilitators made a point to include everyone in discussion and build off people’s suggestions. We also spent time in small groups during the sessions that helped me get to know individual members better.” Another participant praised how their facilitators modeled sharing, “The facilitators asked for input and reflected upon what we shared. The facilitators shared their personal experiences, including areas for improvement, and celebrated the successes of the participants.” Overall, the LC structure was described as a space for discussing and sharing experiences, and facilitators were often credited with creating opportunities for participants to share and connect.

##### Finding Commonalities Amongst Peers

Participants frequently attributed their sense of community to the commonalities and shared sense of community they felt amongst peers. Terms such as “shared goals,” “same issues,” and “common problems” frequently appeared in participants’ open responses. One participant described how much they appreciated a sense of shared objectives amongst peers: “It was great to be together with other professors from STEM fields who were thinking about these [DEI] issues. We came from a shared place of ‘we know we want to do better on this but aren’t sure how’ and were able to openly share our struggles and challenges, including sometimes frustrations with the online course.” Another participant stated that their LC consisted of members from a single professional organization, and in particular, “this was a good way to get to know some of the membership better, and to meet like minded individuals who are invested in DEI and inclusive teaching principles. All members expressed a mutual desire to bring about positive change within our discipline.” Participants also acknowledged that while members of the LC might bring different perspectives, “everyone [who] was there…wanted to learn from each other so we could all do better and be the best version of ourselves in our current and/or future teaching practice.” The shared goal to improve and learn was a key community building characteristic that made participants enjoy these peer interactions.

Feeling a sense of common identities and goals encouraged a sense of continued community for participants. As one participant explained, “I really enjoyed getting to know my community. I think the strongest example of connection was finding someone else who shares a similar history / set of experiences as me prior to college, and learning how this has impacted our teaching in similar ways. We have actually scheduled a follow up meeting this summer together.” Another participant appreciated hearing differing opinions: “Everyone was there with very different experiences with the mindset that they wanted to learn from each other so we could all do better and be the best version of ourselves in our current and/or future teaching practice.” Finding commonality with peers helped participants feel they had a community to help motivate and support them in incorporating inclusive teaching strategies.

##### Feeling Heard & Valued by Peers

Participants most frequently attributed their strong sense of community to the productive interactions they had with their LC peers. They described feeling “heard,” “valued,” and “safe” in the interpersonal dynamics within their facilitated LCs. Participants appreciated the active listening approaches employed by their fellow participants, such as how they “acknowledged” opinions, “carefully listened,” and “showed support and appreciation” during discussions. Importantly, results show that the experiences of participants with marginalized identities (e.g., women and people of color) were not significantly different than their majority peers. For example, one Latina woman described her LC as “an affirming space for individuals with minoritized identities in spite of being a small minority within the group.” Similarly, an East Asian participant shared, “As an international scholar, I had experienced that my voice was not heard and valued in a group discussion because of my English proficiency, different cultural experience, and many other reasons. However, I felt a sense of belonging in my LC because the members of my LC always showed their interests in carefully listening to my opinions and responding to them genuinely.” Participants with minority identities emphasized the sense of safety and respect in the space. As another Latina woman shared, “This is only the second time in my academic career that I have felt that I was in a safe space. I trusted that what we said was kept confidential, which helped us learn from each others’ experiences.” Participants of majority and minority identities credited their colleagues for not only creating a safe space within the LC to engage in highly sensitive discussion topics but also valuing the diverse opinions and experiences brought to the LC space.

### What are the relationships between LC facilitators’ inclusive learning facilitation and participant experiences?

#### Facilitators Created an Inclusive Structure for Participation

LCs necessarily reflect local contexts which differ across institutions, participant audiences, and facilitators. To address whether the Inclusive STEM Teaching Project successfully trained facilitators who are guiding LCs whose participants achieve project-designed learning objectives require exploring activities within each LC. When we compare the participant and facilitator responses for exact and closely aligned questions, we find three distinct domains: (a) overall high congruence, (b) participant agreement is more positive than facilitators, and (c) facilitator agreement is more positive than participants. The first domain displaying the strongest congruence were found in questions addressing LC structure, specifically related to establishing community norms and sharing in the LC (Figure 2).

**Figure 2.**
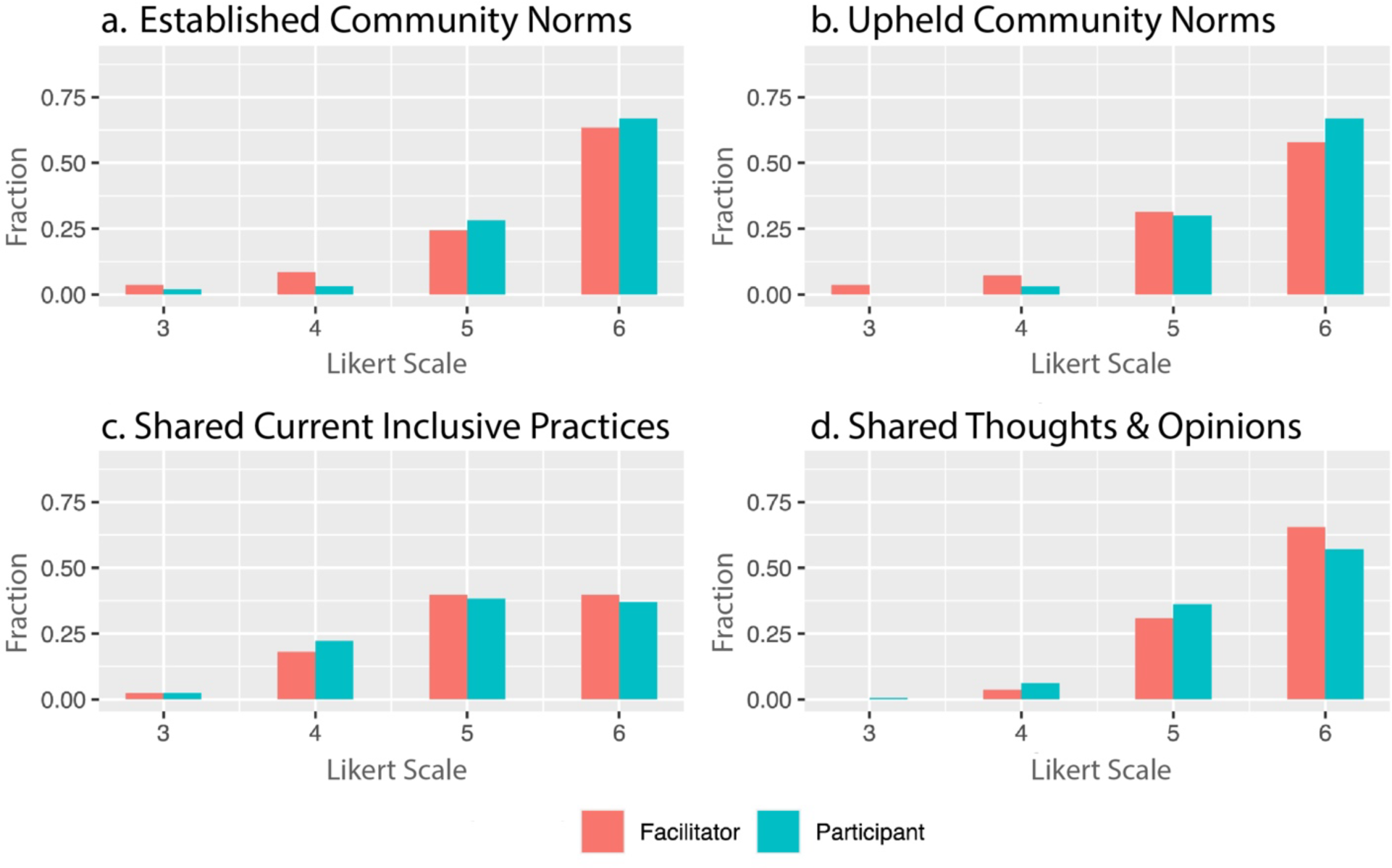
Normalized Fractional Response (0.0 to 1.0) by Cohort of Participant and Facilitator Joint Histograms - High Level of Congruence Between Facilitator and Participant Agreement

Both facilitators and participants agree their LCs established community norms in the first few sessions (Figure 2a and 2b). Participants also self-reported having upheld these community norms and facilitators agreed. One participant with no previous teaching experience specifically described, “When establishing community norms [my facilitators] presented a first draft and had us all actively give feedback and additions to the guidelines.” As a result, the participants felt that the LC facilitators were actively responding to their participants’ feedback and creating a productive learning space. To help confirm that the overall distribution of participant and facilitator responses we plot for these select questions accurately portrays the precise comparison within each LC, we calculated the distribution of the distance between participant and facilitator responses for matching LCs. The question asked about ‘establishing community norms,’ was identical for each population and we find 56% were zero distance or exact matches, 14% (15%) of participants reported one level less (more), with fewer than 10% reporting two or more. In all data where a direct comparison can be made, we find correspondence between matching responses and the overall distributions, lending confidence to our ability to draw conclusions based on overall facilitator and participant responses.

Facilitators and participants also strongly concur that LC participants shared how they were engaging in inclusive teaching practices (Figure 2c) and shared their thoughts and opinions openly (Figure 2d). Hearing peers share ideas was an important part of the LC experience that participants appreciated. Identifying common experiences through sharing fostered participants’ confidence and a willingness to try to implement inclusive practices. In addition, LC participants noted the care with which peers treated each other when sharing questions and experiences.

When asked about the difference between this LC and their past experiences with teaching professional development, one participant stated, “I found the community more open and willing to engage in frank discussions. One person in the group asked a question about using pronouns in an introduction, and I thought everyone involved in the conversation (including the questioner) handled the situation very well.” Another participant emphasized that, “[the] ability to ask questions and get feedback on ideas without worrying about being judged for a misstep - knowing that the feedback and correction would be delivered without judgment for mistakes made.” Facilitator responses paralleled participants’ descriptions of the LC experience. One facilitator stated, “My co-facilitator and I encouraged collaborative problem-solving when a participant voiced a struggle they were having. We also regularly shared [our] own struggles and specific practices we were trying with our own students.” Another facilitator highlighted, “We regularly had a report out of the practices that were introduced in the modules. The facilitators and participants were regularly encouraged to share how they were using inclusive teaching practices and asked to reflect on how this was improving their teaching. The community focused on application of material.” Participants and facilitators both agreed that sharing and discussing opinions and current inclusive practices were significant features of the LC, and in particular participants emphasized feelings of safety and motivation to implement the inclusive teaching practices cultivated by this sharing. Facilitators helped construct this space through setting community norms and structuring peer-sharing focused activities and participants, in turn, self-reported a strong sense of community in the discussion space.

#### Facilitators and Participants Built a Sense of Community

Another group of facilitator and participant comparison histograms visually showed a general agreement with the intentional structured LC environment, but participants were more positive than facilitators (Figure 3). Participants overall highly agreed that facilitators had encouraged participants to guide the direction of community discussions (Figure 3a), were responsive to feedback and made in-time changes to sessions (Figure 3b), and modeled inclusive teaching practices (Figure 3d). Participants also strongly agreed that they felt a sense of community, whereas facilitators expressed less strong agreement that *they created* a sense of community among the participants (Figure 3c), or that they were quite as responsive, or that they were quite as successful in modeling inclusive practices as participants experienced.

**Figure 3.**
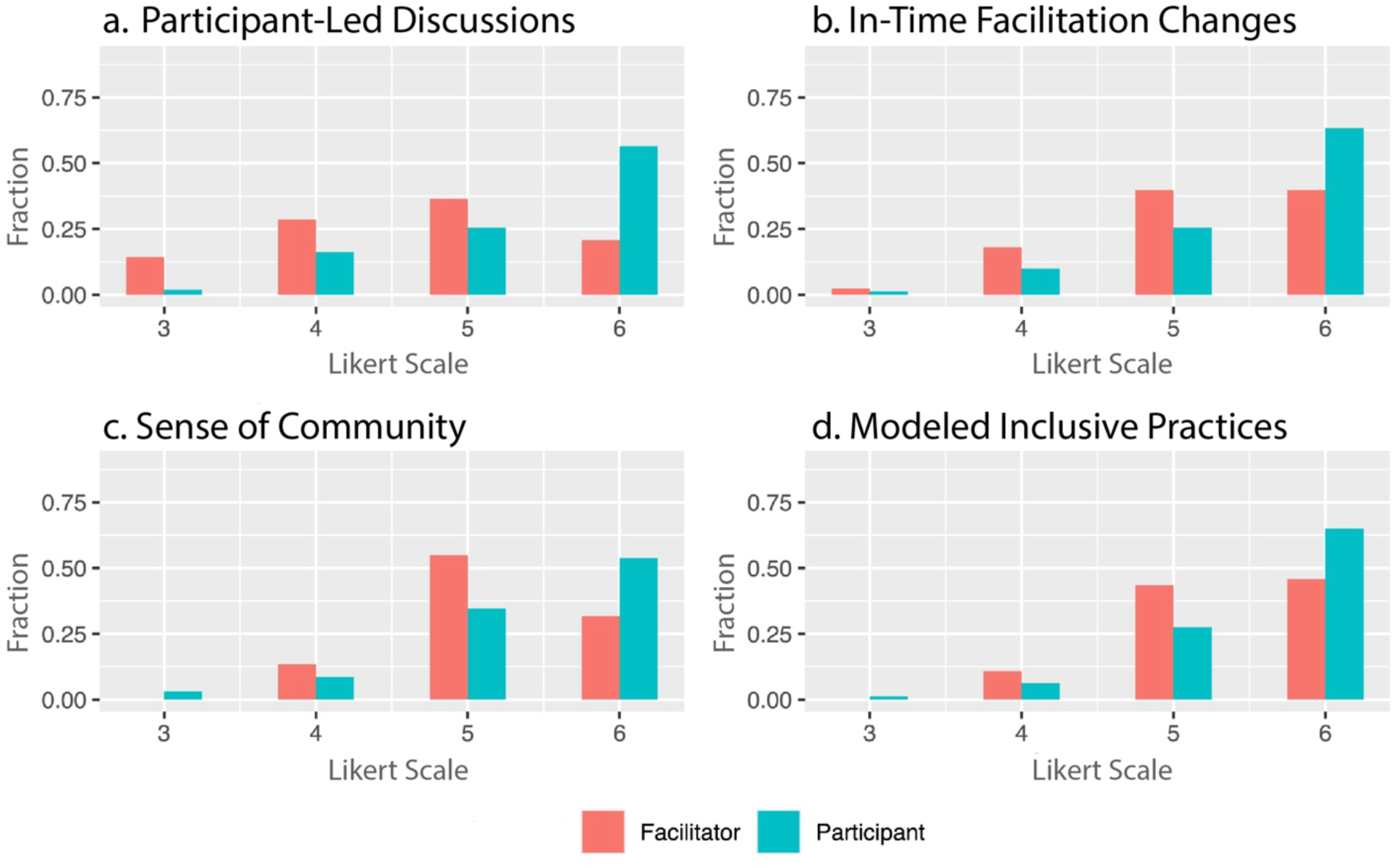
Normalized Fractional Response (0.0 - 1.0) by Cohort of Participant and Facilitator Joint Histograms - Participant responses greater agreement with statement than facilitators.

In addition to the social conforming pressure to under report one’s own positive actions, facilitators may also have felt unsure or less confident that they specifically created the sense of community they envisioned and instead may have felt the participants contributed to this sense of support themselves, consistent with our study of the facilitators (Codding *et al*., 2024). While facilitators often described strategies such as introductions, co-creating community guidelines, and checking-in as methods to create community, participants overwhelmingly credited productive interactions with facilitator and peer actions as the reasons for feeling a sense of community. One participant praised the support from their peers: “I feel that if something does not go well I have people who I can troubleshoot it with. Hearing that others struggle too helps me stay motivated to keep trying things. Hearing about other people’s successes also helps me stay motivated because I would like to have the same success.” Facilitators also emphasize their purposeful desire to create a space in which participants can experience productive interactions. As one facilitator explained, “We came in with a facilitation plan but readily shifted depending on the interests of the group.” Another facilitator explained their approach with even more specificity, “We would regularly expand or reduce the time allotted for activities based on participant discussions or questions. If participants were particularly interested in a topic or challenge, it was easy to expand the time allotted for that activity. If a topic was one that people felt comfortable with and didn’t need as much time for, we would move on or ask burning questions to address what the participants wanted to discuss.”

Many participants echoed facilitators’ intentional approach to providing a general structure to the LC meetings while willingly letting participants take “control” and “steer” conversations towards topics they were interested in exploring. When asked how facilitators were responsive to participant feedback, participants praised facilitators’ willingness to adapt the planned structure of their LCs. One participant pointed out, “[The facilitators] were responsive to the needs of individuals who were struggling with concerns outside of the LC and they adjusted our progression across a session in response to the times when we had a more in-depth conversation and engagement.” Another participant recounted a more significant change in course structure: “One week our material lined up with an on-campus training we’d also all signed up for, so we merged those together instead of having a higher burden for time that week and rehashing the material over and over.” Because facilitators often encouraged participants to speak with each other and responded to fruitful discussions by “stepping back,” this may explain why participants expressed high agreement that they were able to actively lead the LC discussions and felt a sense of community.

Finally, in the third domain we observe that while everyone agrees that participants shared their experiences and engaged in and planned to engage in inclusive teaching practices, participants under report the extent to which they shared or engaged relative to what facilitators reported that they observed among those same participants (see Figure A in Supplemental Material). Under reporting one’s own contributions or expectations would also be typical.

### How are participants planning to implement inclusive teaching practices following LC participation?

#### Planned Changes Reflect ISTP Course Content and LC Activities

The ultimate purpose of the LC and course in inclusive teaching was to encourage learners to implement inclusive instructional practices that support the success of all students. When asked what changes they planned to make to their teaching after participating in the LC, participants provided examples aligned with the breadth and depth of the ISTP online course curriculum and corresponding LC activities. For example, many participants expressed their intent to make important changes to their syllabi. These changes align with an LC activity in which participants bring their syllabi to critically discuss with their peers and identify areas for improvement. Participants specifically noted their intent to use more welcoming language, add a diversity statement, outline course objectives, and provide clearer instructions and expectations regarding class activities such as group work and assignments. Similarly, participants shared that they intend to develop more inclusive assessments and grading practices. One participant discussed their plan to implement these changes by including more non-written assignments and allowing students to propose their own due dates. Another participant described adding an asset-based question to their introductory survey to accommodate students’ diverse learning styles.

Participants also indicated their plans to actively center student identities in their classroom. For example, several participants said they would invite students to share their pronouns at the start of class, again echoing their experience as LC participants. One participant stated, “[I will be] much more mindful of pronouns, gender imbalances in group work, and cognizant of barriers that may be higher [for] some students than others.” Another participant shared they plan to “do more to find ways to bridge cultural gaps between international and domestic students. I’m even more committed to helping first-gen, low income, and racial minority students succeed.” Participants also described supporting student identities by acknowledging and addressing the ways in which STEM fields and higher education have traditionally marginalized underrepresented identities. For example, one participant indicated they would place “more focus on helping students connect their own social identities to what they are studying and learning.”

Participants not only aimed to learn more about their students’ identities, but their responses also indicated their commitment to encouraging a diverse STEM identity and explicitly acknowledging DEI issues such as positionality and privilege in the field. As one participant explained, “I think it is always important to explicitly inform students that the STEM field is enriched by diversity and that everyone’s voice matters. I plan to show students examples of diverse contributors to the STEM field, in the hopes that they will see themselves there.” Many participants planned to incorporate diverse contributors to the STEM field in their curriculum and to explicitly discuss DEI topics in their classrooms: “[I am planning to be] more mindful of the identities, intersectionality, and lived experiences of my students. I hope to make the course relevant to all my students by highlighting many different communities that are affected by and can benefit from earth science education. Specifically, I plan to work to bring more diverse voices into the classroom and share how geosciences understanding and research can be deployed to support environmental and social justice goals.” Most participants were able to describe how they would implement inclusive teaching practices that aligned with ISTP’s modules on student and instructor identity, syllabus design, and active learning strategies.

#### Continuing to Develop as Inclusive Educators

When asked to describe how they would continue their engagement with DEI activities, participants indicated their intent to continue pursuing DEI-focused professional development opportunities. Specifically, participants indicated that they would pursue opportunities for further development through their institution and department in addition to participating in independent activities such as attending workshops, reading more literature, and developing a greater awareness of their own implicit biases outside of the classroom. The majority of participants indicated they would continue to pursue DEI-focused curricular changes—a key focus of the ISTP online course.

## Discussion

In this study conducted over the course of two years and five runs of identity-focused DEI inclusive teaching LCs with an associated online course, we identify three key findings. First, a large-scale comparison of facilitator and participant responses in the same LCs suggest very high fidelity of facilitator’s actual to intended practice resulting in strongly positive and surprisingly congruent participant experiences. Second, we observe virtually no significant differences across race, gender, institutional type or faculty status. Third, we find that while participants acknowledge their facilitator’s inclusive practices, they also credited their fellow participants with helping them develop the confidence to plan and implement inclusive teaching in STEM classrooms.

Our large-scale, direct comparative analysis of the participant and project-trained facilitator experience in higher education LCs across a wide and diverse set of institutional contexts has not yet been researched to our knowledge. Our findings also further current research on effective facilitation techniques in online course-associated LCs (Blum-Smith *et al*., 2021; House *et al*., 2023; McDaniels *et al*., 2016). Through qualitative and quantitative responses, participants were able to describe the ways in which facilitators created LC structures, such as community norms, in-time adjustments to allow for continued discussion, and a facilitated space for sharing experiences. Facilitators likewise confirmed their intentional application of these inclusive facilitation strategies within the LC (see also Codding *et al*., 2024). The level of congruence of facilitation and participant experience is surprising given the different contexts and the challenge of equity-minded work in today’s higher education environment. Participant confirmation that facilitators successfully translated their intent to create a supportive LC structure reinforces the effectiveness of the ISTP effort and of other large-scale train-the-trainer models in inclusive teaching and in STEM mentoring practices (Pfund *et al*., 2017; Rogers *et al*., 2018).

Our findings demonstrate a uniformity of experience. Facilitators were uniformly vetted, trained, collectively supported with synchronous drops-ins that led to consistent outcomes (Codding *et al*., 2024). Though we might have expected factors such as race, gender, institutional context, and faculty status among participants to indicate areas of potential difference of experience, to a large extent they did not. This finding is echoed in the course outcomes of learners in the larger online course itself (Calkins *et al*., 2023), where we have attributed the

uniformity to the self-selection bias of course participants who largely represent a similarly motivated, DEI-experienced, and equity-minded group regardless of demographic of institutional difference. Additionally, in examining three of the five course runs reported here (summer 2021, fall 2021, spring 2022), we found that participating in the online course alone versus participating in the online course and our course-aligned LC did not lead to significantly greater gains on our course learning outcomes scales—a pre-post-course matched survey comparison of awareness, confidence, reflection on and intent to implement inclusive teaching. Although this is consistent with the emphasis on peer engagement we see in data here, it is a bit surprising, nevertheless. LC facilitators have reported anecdotally, and our online course discussion moderators have confirmed, that LC participants often had less online course engagement, perhaps because the LC provided the same peer engagement and support that online-only learners sought through course discussions and affinity groups (Calkins *et al*., 2023).

While participants appreciated how facilitators constructed an inclusive discussion space for them in their LC, they also often credited *motivation from their LC peers* as helping them develop confidence and troubleshoot concepts related to inclusive teaching: “Hearing that others had similar struggles identifying ways to implement inclusive teaching practices gave me confidence to just get started and recognize that I may make mistakes along the way.” Participants consistently described how engaging in productive discomfort (Taylor & Baker, 2019) with one another helped further learning and overcome a fear of “doing it [inclusive teaching] wrong.” LC participants heard from other colleagues who implemented various pedagogical strategies, shared their own approaches, and asked others to help clarify and deepen their knowledge. A combination of sharing and gaining more expertise in the LC made participants feel heard. In addition, participants enjoyed not only sharing identities with peers and common commitments to DEI work but also hearing different experiences and learning from others with different backgrounds. The value of cultural competency and diverse identities in the LC, and engaging with empathy and mutual support (Cole, 2017; Yosso, 2005) allowed LCs to act as a practice space for instructors to take what they learned from ISTP and apply it.

### Limitations

We compared LC facilitator and participant survey responses in our study, but we had two limitations. First, because the LC participant and facilitator surveys were not *prima facie* designed to be a matched study, only a few of the questions were identical, and we could make only visual and not overall numerical correlations between responses in aggregate. As a result, we were limited on certain quantitative analyses. Second, our facilitator survey measured three evaluation points pre-facilitator training, post-facilitator training, and post-LC through Likert scale responses (see Codding *et al*., 2024), but we measured only one evaluation point, post-LC, for participants. While one of our questions asked participants what they *could do* to be more inclusive in their classrooms, and what they plan to do after the LC, these were qualitative responses not used for comparison. For future studies, we plan to include two evaluation points asking participants what they planned to do (pre-LC) regarding inclusive classroom strategies and DEI activities, and what they plan to do after participating in the LC (post-LC).

### Future Applications and Conclusion

The goal of the nationwide LC portion of ISTP is to support facilitators in delivering LCs to locally advance inclusive teaching within their institutional contexts that help participants develop confidence in implementing inclusive teaching strategies in their classrooms. We hope that this study will encourage facilitators, administrators and potential faculty and future faculty participants to not only continue to spread this work through our effective ISTP LC model, but also to seek to influence their local culture around equity and inclusion in higher education. This study only asked one qualitative question about wider changes to DEI work, including institutional changes, and one quantitative question about engaging in DEI conversations with higher ed stakeholders. Participant responses clearly indicated less confidence in implementing such wide institutional changes, and the goal of ISTP and its LCs will ideally be to foster this motivation for institutional change and create communities of support for higher education instructors at various institutions running LCs. This study has shown how a large-scale training approach can foster a strong community of DEI practitioners and learners who grow from direct application of the course materials they learned: engaging in productive discomfort, reflection, and exploring their own identity, positionality, and power in the classroom.

### Accessing Materials

Inclusive STEM Teaching Project (https://www.inclusivestemteaching.org/)

## Footnotes

1 Facilitator data from the first four course runs (*n* = 71) were analyzed and reported on in a previous publication (Codding *et al*., 2024).

