## Supplemental Material for "Community, Belonging, and Peer-Engagement: Participant Experiences Reflect Consistency of Facilitation Practices in Inclusive STEM Teaching Learning Communities"

### Participant Survey

1. Please rate your agreement with the following statements:

| | | Strongly Disagree | Disagree | Somewhat Disagree | Somewhat Agree | Agree | Strongly Agree |
|---|---|---|---|---|---|---|---|
| a. | I attended a majority of the learning-community sessions. | ○ | ○ | ○ | ○ | ○ | ○ |
| b. | My learning community participation was consistent week to week. | ○ | ○ | ○ | ○ | ○ | ○ |
| c. | My learning community established community norms in the first few sessions. | ○ | ○ | ○ | ○ | ○ | ○ |
| d. | I actively upheld our community's norms. | ○ | ○ | ○ | ○ | ○ | ○ |
| e. | I regularly shared my own experiences. | ○ | ○ | ○ | ○ | ○ | ○ |
| f. | Other members of my learning community shared their thoughts and opinions openly during the sessions. | ○ | ○ | ○ | ○ | ○ | ○ |

2. Please rate your agreement with the following statements:

| | | Strongly Disagree | Disagree | Somewhat Disagree | Somewhat Agree | Agree | Strongly Agree |
|---|---|---|---|---|---|---|---|
| a. | My facilitator(s) regularly encouraged learning community participants to guide (or control) the direction of community discussions. | ○ | ○ | ○ | ○ | ○ | ○ |
| b. | I felt that I could lead or guide the direction of the discussion. | ○ | ○ | ○ | ○ | ○ | ○ |
| c. | I felt a sense of community. | ○ | ○ | ○ | ○ | ○ | ○ |
| d. | My facilitator(s) was/were responsive to me and other learning community participants by making just-in-time changes to the sessions based on feedback. | ○ | ○ | ○ | ○ | ○ | ○ |
| e. | My facilitator(s) regularly highlighted and amplified my contributions. | ○ | ○ | ○ | ○ | ○ | ○ |
| f. | My facilitator(s) intentionally modeled inclusive practices in STEM teaching. | ○ | ○ | ○ | ○ | ○ | ○ |
| g. | My facilitator(s) included meta cognitive discussions of the reasons behind modeling inclusive practices in STEM teaching. | ○ | ○ | ○ | ○ | ○ | ○ |
| h. | I regularly shared ways I was already engaging with inclusive teaching practices. | ○ | ○ | ○ | ○ | ○ | ○ |
| i. | I regularly shared ways I could engage with inclusive teaching practices. | ○ | ○ | ○ | ○ | ○ | ○ |
| j. | I felt regularly encouraged to share how I was already or could engage with inclusive teaching practices. | ○ | ○ | ○ | ○ | ○ | ○ |

3. Please provide one or two examples of ways you were already engaging with inclusive teaching practices.

4. Please provide one or two examples of ways you could engage with inclusive teaching practices.

5. Please provide one or two examples of why you felt a sense of community in your learning community.

6. Please provide one or two examples of how your facilitator(s) was/were responsive to your and other learning community participants' feedback.

7. What learning community activities or approaches do you think were particularly engaging for you?

8. What learning community activities or approaches do you think were least engaging for you?

9. Is there anything you wish the learning community had done and would recommend for future Inclusive STEM Teaching learning communities?

10. Prior to participating in this learning community, please indicate in which of the following diversity, equity, and inclusion (DEI) activities or conversations you participated in. Please include both teaching-related and non-teaching related DEI activities (check all that apply):
    - Attended DEI events outside of my local institution.
    - Attended inclusive teaching events offered at my local institution.
    - Attended inclusive teaching events outside of my local institution.
    - Participated in a learning community (on any subject) at my local institution.
    - Participated in a learning community (on any subject) outside of my local institution.
    - Organized DEI or inclusive teaching events offered at my local institution.
    - Organized DEI or inclusive teaching events outside of my local institution.
    - Facilitated DEI or inclusive teaching conversations within my department (e.g., faculty/department meeting, DEI committee).
    - *Other not included above (please specify): [ ]

11. How many years would you say you have been involved in DEI-related (including inclusive teaching) efforts? (*sliding scale from 0-25 years*)

12. Please rate your agreement with the following statements:

| | | Strongly Disagree | Disagree | Somewhat Disagree | Somewhat Agree | Agree | Strongly Agree |
|---|---|---|---|---|---|---|---|
| a. | As a result of my participation in the learning community, I feel ready to implement inclusive teaching practices. | ○ | ○ | ○ | ○ | ○ | ○ |
| b. | The opportunity to hear that other participants faced the same challenges I do enhanced my confidence to apply inclusive teaching practices. | ○ | ○ | ○ | ○ | ○ | ○ |
| c. | Participating in the learning community has enhanced my motivation to apply inclusive teaching practices in my own instruction. | ○ | ○ | ○ | ○ | ○ | ○ |
| d. | Support from other participants in the learning community enhanced my confidence to implement inclusive teaching practices. | ○ | ○ | ○ | ○ | ○ | ○ |
| e. | This learning community was different than my prior experiences in teaching professional development. | ○ | ○ | ○ | ○ | ○ | ○ |

13. Please describe one or more specific instances of participating in the learning community that has enhanced your readiness to implement inclusive teaching practices.

14. Please describe one or more specific instances of how participating in the learning community was different than your prior teaching professional development experiences.

15. Please rate your agreement with the following statements:

| | Strongly Disagree | Disagree | Somewhat Disagree | Somewhat Agree | Agree | Strongly Agree |
|---|---|---|---|---|---|---|
| My learning community centered social identity, as defined by the explicit discussion of instructor and student social identities as they relate to teaching and learning. | ○ | ○ | ○ | ○ | ○ | ○ |

16. How did the centering (or not) of identity impact your learning community experience?

17. What changes are you planning to make in your teaching as a result of participating in your learning community and the course?

18. What are you changing or planning to change in your DEI activities generally as a result of participating in your learning community and the course?

19. At which institution (and which course run) did you participate in an Inclusive STEM Teaching Project learning community?

20. Did you achieve completion for the online course and receive either an edX or Inclusive STEM Teaching Certificate?
    - Yes, I completed the online portion of the course and received a certificate.
    - No, I didn't complete the online portion of the course.

21. Please share any key reasons you can identify for not completing the online portion of the course.

22. Are there steps your learning community or the project could have taken to improve your likelihood of completion?

23. What is your primary role?
    - Faculty member, lecturer, instructor, or adjunct faculty
    - Graduate student
    - Postdoctoral scholar
    - Staff member
    - * Other (please tell us): [ ]

24. Please specify the faculty status that most closely matches your status:
    - Tenured (associate or full professor status)
    - Tenure-track (assistant professor status)
    - Full-time teaching or instructional on a fixed-term, renewable contract
    - Full-time teaching or instructional on a fixed-term, non-renewable contract
    - Part-time teaching or instructional on a fixed-term, non-renewable contract
    - Research faculty on a fixed-term, renewable contract
    - Research faculty on a fixed-term, non-renewable contract
    - * Other (please tell us): [ ]

25. What percent of your time is spent on the following job duties? Please add to 100 percent.
    - Administration [ ] %
    - Research [ ] %
    - Service [ ] %
    - Teaching [ ] %
    - * Other (please tell us): [ ] [ ] %

26. What is your primary disciplinary affiliation?
    - Arts
    - Biological and life sciences
    - Business and management sciences
    - Chemistry
    - Computer, information, and technological sciences
    - Earth, environmental, atmospheric, and ocean sciences
    - Education
    - Engineering
    - Humanities
    - Law
    - Mathematics and statistics
    - Medical sciences
    - Physical sciences
    - Psychology
    - Social, behavioral, and economic sciences (not including psychology)
    - * Other (please tell us): [ ]

27. At what type of institution are you primarily enrolled or employed?
    - Community college / 2-year institution
    - Comprehensive or regional university (e.g., smaller state school, schools that offer mostly bachelor or master's degrees)
    - Liberal arts college
    - Research university
    - Technical college
    - * Other (please tell us): [ ]

28. Is your institution designated / recognized as one of the following categories?
    - Asian American and Pacific Islander Serving Institution (AAPISI)
    - Hispanic Serving Institution (HSI)
    - Historically Black College and University (HBCU)
    - Predominantly White Institution (PWI)
    - Tribal College/University
    - Other Minority Serving Institution (MSI)
    - I am not sure

29. What is your current gender identity? Please select one or use the write-in option.
    - Gender queer or gender non-conforming
    - Man
    - Nonbinary
    - Transgender
    - Woman
    - * I self-describe as [ ]
    - I prefer not to respond.

30. With which racial/ethnicity group(s) do you identify? Please check all that apply.
    - Alaska Native, American Indian, Native American or Indigenous
    - Asian American
    - Black or African American
    - East Asian
    - Latina/o/x or Hispanic
    - Middle Eastern or Northern African
    - Pacific Islander

- South Asian
- Southeast Asian
- White
- Multiracial
- * I self-describe as [ ]
- I prefer not to respond.

### Facilitator Survey

1. How many times have you facilitated an Inclusive STEM Teaching Project (ISTP) learning community?
    - ○ This was my first time
    - ○ This was my second time *
    - ○ Several times or more *

2. * Thank you for facilitating multiple times. We encourage you to fill out this survey, since learning communities are always different, and you are more experienced for this last course run than previous ones. Still, we acknowledge the time to complete this survey, and if you have responded previously and have had a similar experience, we understand.
    - ○ My learning community experience was different, and I'd like to continue
    - ○ My learning community experience wasn't significantly different, and my earlier responses remain accurate. [*exit survey*]

3. Please rate your agreement with the following statements:

| | | Strongly Disagree | Disagree | Somewhat Disagree | Somewhat Agree | Agree | Strongly Agree |
|---|---|---|---|---|---|---|---|
| a. | The majority of learners who agreed to participate in my learning community participated in a majority of the learning-community sessions. | ○ | ○ | ○ | ○ | ○ | ○ |
| b. | My learning community participation was consistent week to week. | ○ | ○ | ○ | ○ | ○ | ○ |
| c. | My learning community participants were highly engaged in our synchronous online video space. | ○ | ○ | ○ | ○ | ○ | ○ |
| d. | My learning community established community norms in the first few sessions. | ○ | ○ | ○ | ○ | ○ | ○ |
| e. | Members of my learning community actively upheld our community’s norms. | ○ | ○ | ○ | ○ | ○ | ○ |
| f. | Learning community participants regularly shared their own experiences. | ○ | ○ | ○ | ○ | ○ | ○ |
| g. | Members of my learning community shared their thoughts and opinions openly during the sessions. | ○ | ○ | ○ | ○ | ○ | ○ |
| h. | My learning community discussed instructor and student social identities when engaging with the content throughout the modules. | ○ | ○ | ○ | ○ | ○ | ○ |

4. Please rate your agreement with the following statements:

| | | Strongly Disagree | Disagree | Somewhat Disagree | Somewhat Agree | Agree | Strongly Agree |
|---|---|---|---|---|---|---|---|
| a. | I regularly encouraged learning community participants to guide (or control) the direction of community discussions. | ○ | ○ | ○ | ○ | ○ | ○ |
| b. | My learning community was nearly always guided by me (or my co-facilitator) | ○ | ○ | ○ | ○ | ○ | ○ |
| c. | I created a sense of community among the participants. | ○ | ○ | ○ | ○ | ○ | ○ |
| d. | I was responsive to learning community participants by making in-time changes to the sessions based on feedback. | ○ | ○ | ○ | ○ | ○ | ○ |

| | | | | | | | |
|---|---|---|---|---|---|---|---|
| e. | My facilitation included regularly highlighting and amplifying the contributions of learning community participants. | ○ | ○ | ○ | ○ | ○ | ○ |
| f. | My facilitation included intentional modeling of inclusive practices in STEM teaching. | ○ | ○ | ○ | ○ | ○ | ○ |
| g. | My facilitation included meta cognitive discussions of the reasons behind modeling inclusive practices in STEM teaching. | ○ | ○ | ○ | ○ | ○ | ○ |
| h. | I regularly shared ways to engage with inclusive teaching practices. | ○ | ○ | ○ | ○ | ○ | ○ |
| i. | I regularly encouraged the learning community participants to share ways one can engage with inclusive teaching practices. | ○ | ○ | ○ | ○ | ○ | ○ |

5. * Please provide one or two examples of how you created a sense of community among participants.

6. ** Please provide one or two examples of how you were responsive to learning community participants by making in-time changes to the sessions based on feedback.

7. *** Please provide one or two examples of how you regularly encouraged learning community participants to engage with inclusive teaching practices.

8. What activities or approaches do you think were particularly engaging for your learning community participants?

9. What activities or approaches do you think were particularly impactful for your learning community participants?

10. Is there anything you would do differently when facilitating a future learning community?

11. What are the most significant 2 or 3 reasons you chose to facilitate an ISTP learning community?

12. Prior to facilitating this learning community, please indicate in which of the following DEI activities or conversations you participated (check all that apply):
    - Attended DEI events offered at my local institution
    - Attended DEI events outside of my local institution
    - Organized DEI events offered at my local institution
    - Organized DEI events outside of my local institution
    - Facilitated DEI conversations within my department (e.g. faculty/dept meeting, DEI committee)
    - Facilitated DEI conversations broadly within my local institution (e.g. DEI workshops, events)
    - Facilitated DEI conversations outside of my local institution
    - Other not included above (please specify): [ ]

13. How many years would you say you have been involved in DEI-related work: (*sliding scale from 0-25*)

14. **Prior to participating** in the ISTP **facilitator training**, how confident did you feel:

| | | Extremely Confident | Very Confident | Confident | Somewhat Confident | A Little Confident | Not at All Confident |
|---|---|---|---|---|---|---|---|
| a. | ...facilitating DEI conversations | ○ | ○ | ○ | ○ | ○ | ○ |
| b. | ...creating open dialogue | ○ | ○ | ○ | ○ | ○ | ○ |
| c. | ...creating opportunities for participants to learn from one another | ○ | ○ | ○ | ○ | ○ | ○ |

| | | | | | | | |
|---|---|---|---|---|---|---|---|
| d. | ...leading conversations centered on identity | ○ | ○ | ○ | ○ | ○ | ○ |
| e. | ...leading discussions with higher ed instructors (faculty member, lecturer, instructor, or adjunct faculty) | ○ | ○ | ○ | ○ | ○ | ○ |
| f. | ...sharing your own personal narrative | ○ | ○ | ○ | ○ | ○ | ○ |
| g. | ...managing difficult moments in DEI conversations | ○ | ○ | ○ | ○ | ○ | ○ |

15. **After participating** in the ISTP **facilitator training**, how confident did you feel:

| | | Extremely Confident | Very Confident | Confident | Somewhat Confident | A Little Confident | Not at All Confident |
|---|---|---|---|---|---|---|---|
| a. | ...facilitating DEI conversations | ○ | ○ | ○ | ○ | ○ | ○ |
| b. | ...creating open dialogue | ○ | ○ | ○ | ○ | ○ | ○ |
| c. | ...creating opportunities for participants to learn from one another | ○ | ○ | ○ | ○ | ○ | ○ |
| d. | ...leading conversations centered on identity | ○ | ○ | ○ | ○ | ○ | ○ |
| e. | ...leading discussions with higher ed instructors (faculty member, lecturer, instructor, or adjunct faculty) | ○ | ○ | ○ | ○ | ○ | ○ |
| f. | ...sharing your own personal narrative | ○ | ○ | ○ | ○ | ○ | ○ |
| g. | ...managing difficult moments in DEI conversations | ○ | ○ | ○ | ○ | ○ | ○ |

16. **After facilitating** an ISTP **learning community**, how confident did you feel:

| | | Extremely Confident | Very Confident | Confident | Somewhat Confident | A Little Confident | Not at All Confident |
|---|---|---|---|---|---|---|---|
| a. | ...facilitating DEI conversations | ○ | ○ | ○ | ○ | ○ | ○ |
| b. | ...creating open dialogue | ○ | ○ | ○ | ○ | ○ | ○ |
| c. | ...creating opportunities for participants to learn from one another | ○ | ○ | ○ | ○ | ○ | ○ |
| d. | ...leading conversations centered on identity | ○ | ○ | ○ | ○ | ○ | ○ |
| e. | ...leading discussions with higher ed instructors (faculty member, lecturer, instructor, or adjunct faculty) | ○ | ○ | ○ | ○ | ○ | ○ |
| f. | ...sharing your own personal narrative | ○ | ○ | ○ | ○ | ○ | ○ |
| g. | ...managing difficult moments in DEI conversations | ○ | ○ | ○ | ○ | ○ | ○ |

17. Have you facilitated non-DEI-related learning communities previously?
    - ○ No
    - ○ Yes *

18. * Please explain what, if anything, you found different in facilitating a DEI-related learning community compared to facilitating non-DEI-related learning communities. Please give specific examples.

19. Do you plan to participate in professional development opportunities in the next year to enhance your own facilitation skills?
    - No
    - Yes *

20. * Please describe your plans to participate in professional development opportunities in the next year to enhance your facilitation skills.

21. Did you participate in any of our three learning community facilitator reflection sessions during the course run, held on alternate Wednesdays?
    - Yes, one session **
    - Yes, more than one session **
    - No, I didn't attend *

22. * What prevented your participation?

23. ** How useful were community facilitator reflection sessions?

| Extremely Useless | Useless | Slightly Useless | Slightly Useful | Moderately Useful | Extremely Useful |
|---|---|---|---|---|---|
| ○ | ○ | ○ | ○ | ○† | ○† |

24. † What aspects were most valuable?

25. ** What was missing that you would have found helpful/useful/valuable?

26. Did your learning community offer participants incentives for either participation and/or completion?
    - Yes, we offered incentives *
    - No, we did not offer incentives

27. * Please explain how your learning community utilized incentives, and whether you thought they were effective.

28. In planning or facilitating your learning community, to what extent did you use the following aspects of the facilitator toolkit?

| | | None at All | A Little | Somewhat | Moderately | Very | Extremely |
|---|---|---|---|---|---|---|---|
| a. | ...facilitating DEI conversations | ○ | ○ | ○ | ○ | ○* | ○* |
| b. | ...creating open dialogue | ○ | ○ | ○ | ○ | ○** | ○** |
| c. | ...creating opportunities for participants to learn from one another | ○ | ○ | ○ | ○ | ○*** | ○*** |

29. * How did the facilitator workbook support your learning community?

30. ** How did the email templates support your learning community?

31. *** How did the flyer templates support your learning community?

32. What changes/additions would you recommend for the tools provided?

33. Based on your experience facilitating your learning community this summer, are there other topics and/or skills you would recommend to include in future facilitator training?

34. In what other ways, if any, could the Project support your future facilitation of an ISTP learning community?

35. Please share 1-2 examples of what changes your learning community participants are planning to make in their teaching as a result of participating in your learning community and the course?

36. Please share 1-2 examples of what your learning community participants are changing or planning to change in their DEI activities generally as a result of participating in your learning community and the course?

37. Please share 1-2 examples of what changes you are planning to make in your teaching as a result of facilitating this learning community?

38. Please share 1-2 examples of what you are changing or planning to change in your DEI activities generally as a result of facilitating this learning community?

39. Are you a member of the ISTP, supported through one of the five collaborating institutions (*names redacted*)?
    - ○ No
    - ○ Yes

40. At which institution (and which course run) did you facilitate an ISTP learning community? (*drop down menu*)*

41. * Since you wrote 'Other' please briefly describe your learning community and let us know where it was held.

42. What is your primary role?
    - ○ Faculty member, lecturer, instructor, or adjunct faculty *
    - ○ Graduate student
    - ○ Postdoctoral fellow
    - ○ Staff member
    - ○ Other (please tell us): [ ]

43. * What percent of your time is spent on the following job duties? Please add to 100 percent.
    - ○ Administration [ ] %
    - ○ Research [ ] %
    - ○ Service [ ] %
    - ○ Teaching [ ] %
    - ○ Other (please tell us) [ ] [ ] %

44. What is your primary disciplinary affiliation?
    - ○ Agriculture and natural resource sciences
    - ○ Arts
    - ○ Biological and life sciences
    - ○ Chemistry
    - ○ Computer, information, and technological sciences
    - ○ Earth, environmental, atmospheric, and ocean sciences
    - ○ Education
    - ○ Engineering
    - ○ Humanities
    - ○ Law
    - ○ Mathematics and statistics

- ○ Medical sciences
- ○ Physical sciences
- ○ Psychology
- ○ Social, behavioral, and economic sciences (not including psychology)
- ○ Other (please tell us): [ ]

45. At what type of institution are you primarily enrolled or employed?
    - ○ Community college / 2-year institution
    - ○ Comprehensive or regional university (e.g., smaller state school, schools that offer mostly bachelor or master's degrees)
    - ○ Liberal arts college
    - ○ Research university
    - ○ Technical college
    - ○ Other (please tell us): [ ]

46. Is your institution designated / recognized as one of the following categories?
    - Asian American and Pacific Islander Serving Institution (AAPISI)
    - Hispanic Serving Institution (HSI)
    - Historically Black College and University (HBCU)
    - Predominantly White Institution (PWI)
    - Tribal College/University
    - Other Minority Serving Institution (MSI)
    - I am not sure

47. What is your current gender identity? Please select one or use the write-in option.
    - ○ Gender queer or gender non-conforming
    - ○ Man
    - ○ Nonbinary
    - ○ Transgender
    - ○ Woman
    - ○ I self-describe as: [ ]
    - ○ I prefer not to respond

48. With which racial/ethnicity group(s) do you identify? Please check all that apply.
    - Alaska Native, American Indian, Native American or Indigenous
    - Asian American
    - Black or African American
    - East Asian
    - Latina/o/x or Hispanic
    - Middle Eastern or Northern African
    - Pacific Islander
    - South Asian
    - Southeast Asian
    - White
    - Multiracial
    - I self-describe as: [ ]
    - I prefer not to respond

**Table A.** Matching Participant and Facilitator Survey Questions

| Participant Survey Question | Facilitator Survey Question | Figure |
|---|---|---|
| (1c) My learning community established community norms in the first few sessions. | (3d) My learning community established community norms in the first few sessions. | 2a |
| (1d) I actively upheld our community norms. | (3e) Members of my learning community actively upheld our community's norms. | 2b |
| (2h) I regularly shared ways I was already engaging in inclusive teaching practices. | (4h) I regularly shared ways to engage with inclusive teaching practices. | 2c |
| (1f) Other members of my learning community shared their thoughts and opinions openly during the sessions. | (3g) Members of my learning community shared their thoughts and opinions openly during the sessions. | 2d |
| (2a) My facilitator(s) regularly encouraged learning community participants to guide (or control) the direction of community discussions. | (4a) I regularly encouraged learning community participants to guide (or control) the direction of community discussions. | 3a |
| (2d) My facilitator(s) was/were responsive to me and other learning community participants by making just-in-time changes to the sessions based on feedback. | (4d) I was responsive to learning community participants by making in-time changes to the sessions based on feedback. | 3b |
| (2c) I felt a sense of community. | (4c) I created a sense of community among the participants. | 3c |
| (2f) My facilitator(s) intentionally modeled inclusive practices in STEM teaching. | (4f) My facilitation included intentional modeling of inclusive practices in STEM teaching. | 3d |
| (1e) I regularly shared my own experiences. | (3f) Learning community participants regularly shared their own experiences. | 4a |
| (2i) I regularly shared ways I could engage with inclusive teaching practices. | (4i) I regularly encouraged the learning community participants to share ways one can engage with inclusive teaching practices. | 4b |
| (15) My learning community centered social identity, as defined by the explicit discussion of instructor and student social identities as they relate to teaching and learning. | (3h) My learning community discussed instructor and student social identities when engaging with the content throughout the modules. | 4c |

### Qualitative Codebook

| Parent Code | Child Code | Data Example |
|---|---|---|
| **A. Identity and Awareness** (Instructor and Facilitator focused)<br><br>Ways in which instructors and facilitators use, discuss, value and acknowledge their own and students' identities, as well as identity-based historical inequities in learning environments | **A1. Student/Participant Identity**<br><br>Instructors/Facilitators invite or asks students/participants to share their names, pronouns, and parts of identity to create inclusive learning space through a variety of means (e.g., introductions on the first days, pre-course surveys) | "We asked everyone to create name tags and make them visible during sessions, and had people routinely share something about their current teaching during each session." |
| | **A2. Instructor/Facilitator Identity**<br><br>Instructor/facilitator intentionally shares parts of their personal identity with students/participants | "[I plan to] add a little bit more about me at the beginning of the semester, about myself and my failures." |
| | **A3. Cultivating Diverse STEM Identity**<br><br>Instructor/facilitator incorporates diverse identities and representation in the STEM curriculum (e.g., marginalized individuals' scientific contributions, diverse identities in videos) | "[Participants in the future plan to] Incorporate their identity in introductions, changing readings/videos for representation" |
| | **A4. Acknowledging positionality and privilege**<br><br>Instructors/facilitators describe how their identity relates to privilege and positionality in the role as an instructor in their classrooms. Includes discussions of (in)equity and identity in the field of STEM (e.g., racism, sexism). | "DEI is sensitive subject matter and requires training to be appropriate, sensitive, aware and non-bias. I am still learning about my own bias and unconscious behaviors that may impact others and I am unaware of them. " |
| **B. Inclusive Community**<br><br>Facilitators/Participants describe techniques used to build community and a sense of belonging. Inclusive of future applications. | **B1. Introductions**<br><br>Facilitators/Instructors use the start of class/LC to build community through ice breakers, welcome prompts, and gauging participants. | "[I created a sense of community through] check in slide, icebreakers, welcoming participants, 'hangout time'--> show personal interest and support of participants" |
| | **B2. Community Standards**<br><br>Facilitators/Instructors introduce community guidelines (or build with participants) to establish expectations of conduct and engagement in classroom/LC | "In the first week we went over the community guidelines, and spent some time having the participants generate their questions and ideas for 1) what we wanted the guidelines to be in our space, and 2) how they can generate and invite similar guidelines in their own teaching contexts." |
| | **B3. Welcoming Diverse Forms of Engagement**<br><br>Facilitators/Instructors encourage and acknowledge active participation (e.g., invite participation and thank participants for sharing). Can include UDL techniques, such as captioning videos | "We always encourage everyone to speak up and share, especially those we don't often hear from, and at the time we remind them if they don't want to do it, that is ok too." |
| | **B4. Sharing own Experiences with S/P**<br><br>Instructor/Facilitator share their personal experiences with students/participants to express support in inclusive teaching efforts and encourage community building and discussion | " [I encouraged participants to engage in inclusive teaching by] Providing them with regularly encountered problems most of us face while teaching at Utica and then asking them to apply what they are learning about practices. |

| | | |
|---|---|---|
| | | Recounting problems I have encountered while teaching and how I have solved or intend to solve them.” |
| | **B5. Adapting Course Structure**<br><br>Instructors/Facilitators adjust the course activities, structure, or curriculum to accommodate participant/student needs, feedback, interests, and requests. Includes collecting data/feedback on course climate. | “My co-facilitator did a great job of modeling [responsiveness to learners] - when a short amount of time was left, he polled participants on how to use the remaining time. I was able to use this technique as well in a subsequent session.” |
| **C. LC Group Dynamics**<br><br>Facilitators/Participants reflect on or describe engagement and interactions between group members | **C1. Productive Interactions**<br><br>Facilitators/Participants describe interactions within the LC (e.g., sharing interests, respectfully disagreeing) | “Our community was very friendly, and we all had the space to share (or not) with respect and curiosity.” |
| | **C2. Collaborative Problem-Solving**<br><br>Participants discuss personal challenges and/or engage in collaborative problem-solving. Facilitators encourage participants to collaboratively discuss or problem solve together. | “[I regularly encouraged participants] through the prompts and in any challenges posed by the group as a brainstorm for the group to discuss” |
| | **C3. Highlighting Shared Experiences and Commonalities**<br><br>Facilitators/Participants identify ways in which shared experiences or commonalities impact LC | “I would intentionally refer to their unique shared experiences of training K-12 teachers and as well as their investment in the future of inclusive teaching practices.” |
| | **C4. Continuing Community Beyond LC**<br><br>Participants note an interest in continuing their LC communities after the course | “We should have exchanged information to keep in touch beyond the course.” |
| **D. Discussion Approaches**<br><br>Ways in which facilitators organized discussion opportunities and ways participants/instructors describe using discussion approaches in their classrooms now or in the future | **D1. Large Group Discussion**<br><br>Participants or Facilitators describe large group discussion or interacting with fellow participants as productive to learning (they do not include describing it as problem-solving or community building) | “I think the large group discussions that allowed for the organic revealing of insights or pointed teachable moments, from the co-facilitators were particularly impactful with this group.” |
| | **D2. Small Group Work/Discussions**<br><br>Participants or Facilitators describe the use of small group discussions that then typically report back to the larger group | “I think they really like the small group discussions because they were able to exchange experiences and practices as well as build their network. I also think that liked watching and discussing the Kels video.” |
| | **D3. Participant-lead discussions**<br><br>Facilitators encourage participants to be co-facilitators or lead the discussion topics and length of time discussing topics. | “We rotated facilitating discussions--and community members volunteered to co-facilitate”’ |
| | **D4. Reflection**<br><br>Facilitators encourage participants to reflect on their current and future application of inclusive teaching practices, includes metacognitive reflection | “We regularly had a report out of the practices that were introduced in the modules. The facilitators and participants were regularly encouraged to share how they were using inclusive teaching practices and asked to reflect on how this was |

| | | |
|---|---|---|
| | | improving their teaching. The community focused on application of material.” |
| | **D5. Multi-Modal Discussions**<br><br>Using technology to allow for different methods of engagement/discussion | “We used the Yammer app from Microsoft to create a discussion group. Participants posted a favorite, surprising, or thought-provoking line from the ISTP Module on Yammer each week. Folks from the two different meeting times could see and reply to each other's posts.” |
| **E. Teaching & Pedagogy**<br><br>Various inclusive, active learning strategies and activities described by instructors/facilitators or participants. | **E1. Increased Transparency & Inclusive Language**<br><br>Facilitators/Instructors increase transparency of pedagogy and instruction (e.g., explicitly explaining the “why” of an assignment, improving instructions, expounding on course policies in syllabus, identifying learning outcomes) | “[Learning participants plan to] "Include more description of identity in their introduction to the course. Remove language from the syllabus that might be non-inclusive." |
| | **E2. Active Learning**<br><br>Facilitators and Instructors describe active learning techniques (e.g., minute papers, clickers, jigsaw) | “Deemphasizing lecturing, flipped classroom, problem-based learning.” |
| | **E3. Providing/Curating Resources**<br><br>Facilitators/Instructors describe providing additional resources that help support learning and learners (e.g., shared links and notes, resource banks, curated lists). | “We created [our] own listserv and [shared] resources among ourselves.” |
| | **E4. Inclusive Assessment & Grading**<br><br>Instructors describe using assessment or grading strategies that are more inclusive of a variety of learners | “I have revamped my grading system to be standards-based and have communicated course objectives to students more clearly than in the past. Policy changes include revisions and more flexible timelines for learning. I have looked carefully at how assignments are presented and phrased (for my non-native speakers) and invited students to share information about identities, if they choose. I plan to include an anonymous "reporting tool" on my course webpage so students can communicate issues as they arise.” |
| | **E5. Concrete Application**<br><br>Encouraging applications of inclusive teaching practices (or course content) outside the classrooms. May include “homework” | “Modeling techniques during our sessions was one key way we did that, and setting 'homework' for participants to try new things in between our sessions was another.” |
| | **E6. Soliciting Feedback**<br><br>Facilitator/Instructor solicits feedback from participants/students on the course/LC | “Finding ways to restructure feedback to be more responsive to students’ needs.” |

**Table B.** Likert Scale Responses from LC Participants

| | Strongly Disagree | Disagree | Somewhat Disagree | Somewhat Agree | Agree | Strongly Agree | Mean | SD |
|---|---|---|---|---|---|---|---|---|
| I attended a majority of the learning-community sessions | 1 (0.61%) | 1 (0.61%) | 3 (1.82%) | 12 (7.27%) | 32 (19.39%) | 116 (70.30%) | 5.55 | 0.84 |
| My learning community participation was consistent week to week | 2 (1.21%) | 5 (3.03%) | 10 (6.06%) | 17 (10.30%) | 42 (25.45%) | 89 (53.94%) | 5.18 | 1.15 |
| My learning community established community norms in the first few sessions | 3 (1.83%) | 1 (0.61%) | 3 (1.83%) | 5 (3.05%) | 45 (27.44%) | 107 (65.24%) | 5.49 | 0.93 |
| I actively upheld our community's norms | 2 (1.21%) | 0 (0.0%) | 0 (0.0%) | 5 (3.03%) | 49 (29.70%) | 109 (66.06%) | 5.58 | 0.74 |
| I regularly shared my own experiences | 1 (0.61%) | 0 (0.0%) | 3 (1.82%) | 22 (13.33%) | 56 (33.94%) | 83 (50.30%) | 5.31 | 0.85 |
| Other members of my learning community shared their thoughts and opinions openly during the sessions | 1 (0.61%) | 1 (0.61%) | 1 (0.61%) | 10 (6.06%) | 59 (35.76%) | 93 (56.36%) | 5.45 | 0.78 |
| My facilitator(s) regularly encouraged learning community participants to guide (or control) the direction of community discussions | 1 (0.61%) | 2 (1.22%) | 3 (1.83%) | 26 (15.85%) | 41 (25%) | 91 (55.49%) | 5.30 | 0.95 |
| I felt that I could lead or guide the direction of the discussion | 2 (1.22%) | 3 (1.83%) | 9 (5.49%) | 55 (33.54%) | 58 (35.37%) | 37 (22.56%) | 4.68 | 1.02 |
| I felt a sense of community | 1 (0.61%) | 2 (1.21%) | 5 (3.03%) | 14 (8.48%) | 56 (33.94%) | 87 (52.73%) | 5.32 | 0.92 |
| My facilitator(s) was/were responsive to me and other learning community participants by making just-in-time changes to the sessions based on feedback | 1 (0.61%) | 2(1.22%) | 2 (1.22%) | 16 (9.76%) | 41 (25%) | 102 (62.20%) | 5.44 | 0.89 |
| My facilitator(s) regularly highlighted and amplified my contributions | 1 (0.61%) | 0 (0.0%) | 4 (2.44%) | 22 (13.41%) | 63 (38.41%) | 74 (45.12%) | 5.24 | 0.85 |
| My facilitator(s) intentionally modeled inclusive practices in STEM teaching | 1 (0.61%) | 2 (1.23%) | 2 (1.23%) | 10 (6.13%) | 44 (26.99%) | 104 (63.80%) | 5.49 | 0.85 |
| My facilitator(s) included meta cognitive discussions of the reasons behind modeling inclusive practices in STEM teaching | 3 (1.84%) | 1 (0.61%) | 8 (4.91%) | 24 (14.72%) | 59 (36.20%) | 68 (41.72%) | 5.08 | 1.05 |
| I regularly shared ways I was already engaging with inclusive teaching practices | 1 (0.61%) | 1 (0.61%) | 4 (2.44%) | 36 (21.95%) | 62 (37.80%) | 60 (36.59%) | 5.05 | 0.92 |

| | | | | | | | | |
|---|---|---|---|---|---|---|---|---|
| I regularly shared ways I could engage with inclusive teaching practices | 1 (0.61%) | 0 (0.0%) | 4 (2.42%) | 32 (19.39%) | 68 (41.21%) | 60 (36.36%) | 5.10 | 0.86 |
| I felt regularly encouraged to share how I was already or could engage with inclusive teaching practices | 1 (0.61%) | 1 (0.61%) | 3 (1.82%) | 17 (10.30%) | 55 (33.33%) | 88 (53.33%) | 5.35 | 0.86 |
| As a result of my participation in the learning community, I feel ready to implement inclusive teaching practices | 1 (0.61%) | 2 (1.21%) | 2 (1.21%) | 34 (20.61%) | 81 (49.09%) | 45 (27.27%) | 4.98 | 1.15 |
| The opportunity to hear that other participants faced the same challenges I do enhanced my confidence to apply inclusive teaching practices | 1 (0.61%) | 1 (0.61%) | 3 (1.82%) | 13 (7.88%) | 52 (31.52%) | 95 (57.58%) | 5.41 | 0.93 |
| Participating in the learning community has enhanced my motivation to apply inclusive teaching practices in my own instruction | 1 (0.61%) | 1 (0.61%) | 1 (0.61%) | 17 (10.30%) | 47 (28.48%) | 98 (59.39%) | 5.44 | 0.74 |
| Support from other participants in the learning community enhanced my confidence to implement inclusive teaching practices | 1 (0.61%) | 2 (1.21%) | 3 (1.82%) | 19 (11.52%) | 61 (36.97%) | 79 (47.88%) | 5.29 | 0.85 |
| This learning community was different than my prior experiences in teaching professional development | 1 (0.61%) | 6 (3.66%) | 12 (7.32%) | 31 (18.90%) | 54 (32.93%) | 60 (36.59%) | 4.92 | 0.78 |
| My learning community centered social identity, as defined by the explicit discussion of instructor and student social identities as they relate to teaching and learning | 1 (0.62%) | 3 (1.85%) | 4 (2.47%) | 29 (17.90%) | 69 (42.59%) | 56 (34.57%) | 5.05 | 0.92 |

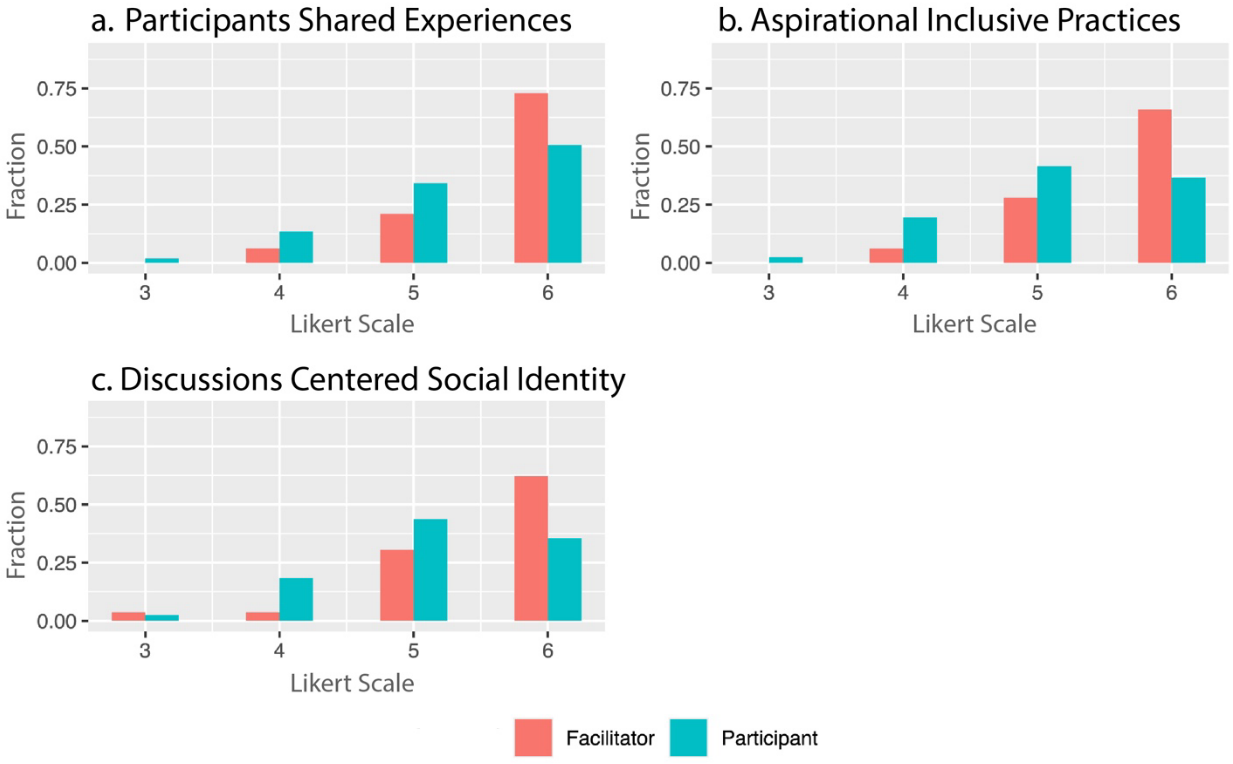


**Figure A.** Normalized fractional response by cohort of participant and facilitator joint histograms - participants' responses less agreement with statement than facilitators.
